# *In vitro* reconstitution of branched microtubule nucleation

**DOI:** 10.1101/700419

**Authors:** Ammarah Tariq, Lucy Green, J Charles G. Jeynes, Christian Soeller, James G. Wakefield

## Abstract

Eukaryotic cell division requires the mitotic spindle, a microtubule (MT)-based structure which accurately aligns and segregates duplicated chromosomes. The dynamics of spindle formation are determined primarily by correctly localising the MT nucleator, *γ*-<u>Tu</u>bulin <u>R</u>ing <u>C</u>omplex (*γ*-TuRC)^1-4^, within the cell. A conserved MT-associated protein complex, Augmin, recruits *γ*-TuRC to pre-existing spindle MTs, amplifying their number, in an essential cellular phenomenon termed “branched” MT nucleation^5-9^. Here, we purify endogenous, GFP-tagged Augmin and *γ*-TuRC from *Drosophila* embryos to near homogeneity using a novel one-step affinity technique. We demonstrate that, *in vitro*, while Augmin alone does not affect Tubulin polymerisation dynamics, it stimulates *γ*-TuRC-dependent MT nucleation in a cell cycle-dependent manner. We also assemble and visualise the MT-Augmin-*γ*-TuRC-MT junction using light microscopy. Our work therefore conclusively reconstitutes branched MT nucleation. It also provides a powerful synthetic approach with which to investigate the emergence of cellular phenomena, such as mitotic spindle formation, from component parts.

---

Branched MT nucleation, and its dependence on Augmin and *γ*-TuRC, generates the bulk of MTs required for both meiotic and mitotic spindle formation^10,11^ and has been visualised *in vivo* in *Drosophila, Xenopus*, plants, and humans^5,6,9-14^. However, understanding, and *in vitro* reconstitution of, this phenomenon has been hampered by methodological constraints relating to purification of functional protein complexes^15,16^; Augmin is composed of 8 subunits, while the *γ*-TuRC is a ~2MD protein complex containing multiple copies of at least 5 proteins, including 14 molecules of *γ*-Tubulin^1-4,17^. *In vitro* studies generally use proteins that have been individually- or co-expressed and purified in heterologous systems^18,19^, where folding and post translational modifications crucial to function may not occur. Although purification of protein complexes from autogenous cells can be achieved using affinity-based methods, non-specific binding of contaminating proteins and difficulties in releasing purified proteins from affinity matrices are major problems.

We therefore developed an approach to allow the isolation of intact, functional Augmin and *γ*-TuRC, to test the hypothesis that these two complexes are necessary and sufficient for branched MT nucleation. The approach is based on biotinylated, amine-reactive thiol- or photo-cleavable linkers, Sulfo-NHS-SS-Biotin and PC-Biotin-NHS (**Figure 1a**). Stepwise incubation of the ~12kD camelid anti-GFP nanobody, GFP-TRAP, with either of these linkers resulted in covalent linkage, while subsequent incubation with a Streptavidin Agarose matrix led to stable tri-partite reagents – GFP-TRAP-Sulfo beads and GFP-TRAP-PC beads; where GFP-TRAP is immobilised, but cleavable through the addition of DTT or exposure to UV light, respectively (**Figure 1a**).

**Figure 1.**
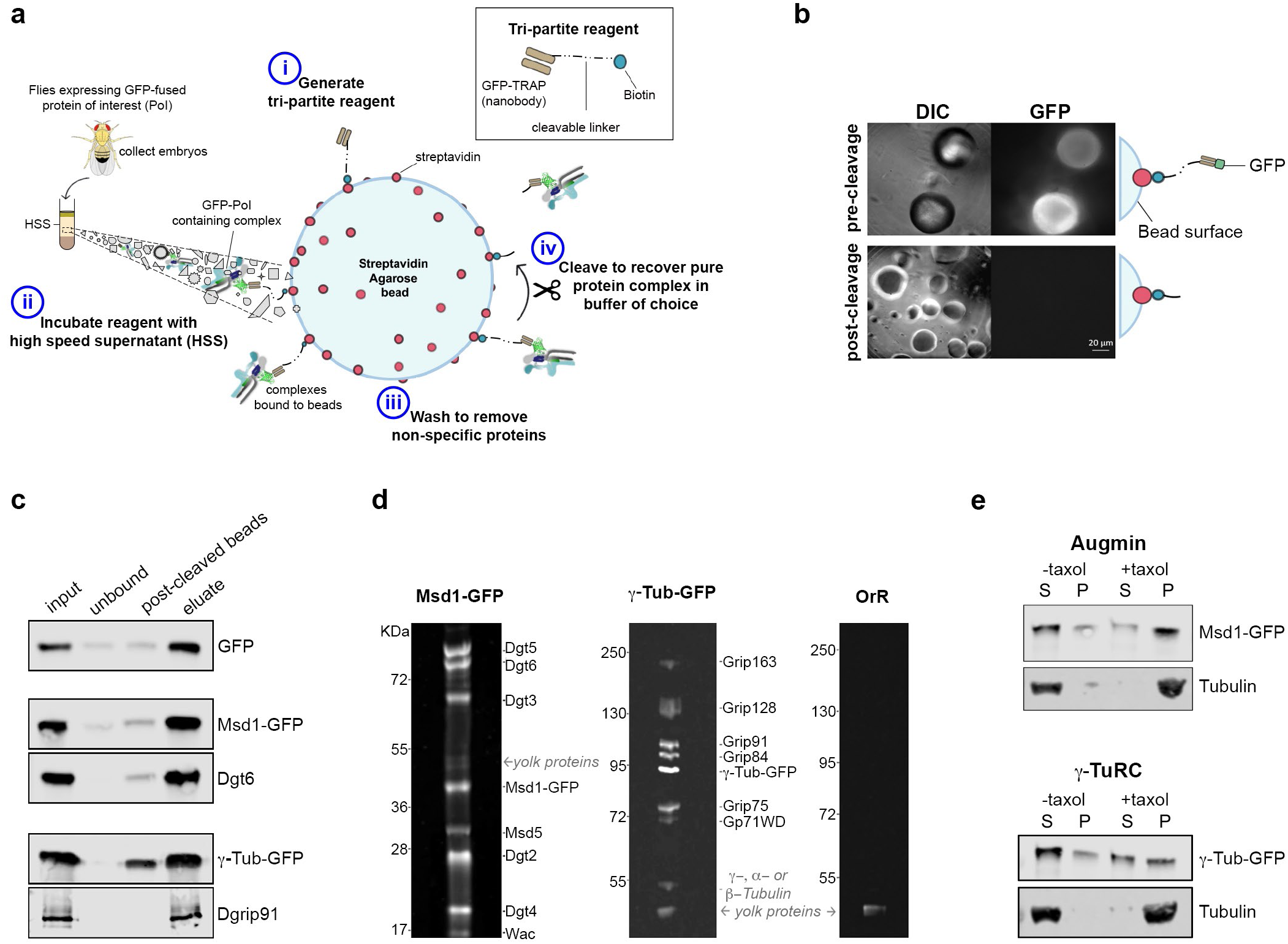
Isolation of functional γ-TuRC and Augmin using cleavable affinity purification. **a**. sketch of purification methodology. **b**. images of GFP-TRAP-Sulfo beads after incubation with GFP, pre- and post-cleavage by 50mM OTT. c. Western blots demonstrating the isolation and cleavage of proteins of interest and interactors. GFP, Msd1-GFP and γ-Tubulin-GFP, present in embryo extracts (input) are efficiently depleted upon incubation with GFP-TRAP-Sulfo beads (unbound), released in 50mM OTT (post-cleaved beads) and present in the eluate. Ogt6, a subunit of Augmin, co-elutes with Msd1-GFP while Ogrip91, a subunit of γ-TuRC, co-elutes with γ-Tubulin-GFP. **d**. SYPRO-ruby stained gels of post - cleaved eluates from control embryos (OrR), or embryos expressing the Augmin subunit Msd1-GFP or γ-Tubulin-GFP. **e**. Western blots of *in vitro* MT co-sedimentation assays using GFP-TRAP-Sulfo isolated complexes. In the absence of taxol (-), Tubulin, pure Augmin (Msd1-GFP) and pure γ-TuRC (γ-Tubulin GFP) remain in the supernatant (S). In the presence of taxol (+), Tubulin polymerises and is present in the pellet (P). Augmin and γ-TuRC co-sediment.

To test these reagents, beads were incubated with bacterially expressed and purified 6xHis-GFP, and extensively washed. Individual beads fluoresced with varying intensity and, upon brief exposure to 50 mM DTT or UV light, fluorescence decreased, concomitant with an increase in the surrounding medium (**Figure 1b**; Supplementary Figure 1a,b). Western blot analysis confirmed >90% and ~60% of GFP was released from GFP-TRAP-Sulfo and GFP-TRAP-PC beads, respectively, following cleavage (**Figure 1c**; Supplementary Figure 1c).

We next sought to determine whether this “<u>cl</u>eavable <u>A</u>ffinity <u>P</u>urification” (cl-AP) could be used to isolate Augmin and *γ*-TuRC. We have previously shown that *Drosophila* Augmin can be purified from extracts of early embryos expressing a GFP-tagged variant of the Msd1 subunit^20^. *Drosophila* embryos have also been used to purify the *γ*-TuRC^2,17^, and flies expressing *γ*-Tubulin-GFP are available^21^. As branched MT nucleation is essential during mitosis, we used embryos arrested in a metaphase-like state through incubation with the proteasomal inhibitor, MG132^22^. Both Msd1-GFP and *γ*-Tubulin-GFP were efficiently immobilised on GFP-TRAP-Sulfo or GFP-TRAP-PC beads and western blotting confirmed that, upon cleaving, Msd1-GFP and *γ*-Tubulin-GFP were concentrated in the eluate, with other subunits of the complexes co-eluting (**Figure 1c**; Supplementary Figure 1d). To test the purity of the complexes, we subjected MG132-treated control, Msd1-GFP or *γ*-Tubulin-GFP embryo extracts to GFP-TRAP-Sulfo cl-AP followed by gel electrophoresis and SYPRO-ruby staining of eluates (**Figure 1d**). Bands corresponding to each subunit of both Augmin and *γ*-TuRC were identified at intensities expected for the known stoichiometric relationships between subunits^17^. One additional set of low intensity bands was seen in all eluates, at ~45kD; almost certainly corresponding to yolk proteins - the most abundant proteins in *Drosophila* early embryos^23^.

Both *γ*-TuRC and Augmin bind MTs in co-sedimentation assays^5, 24-25^. We therefore incubated mitotic Augmin-GFP or *γ*-TuRC-GFP with pure Tubulin, in the presence of GTP and taxol to promote MT polymerisation, sedimenting through a glycerol cushion to separate MTs and MT associated proteins from soluble Tubulin and non-MT binding proteins (**Figure 1e;** Supplementary Figure 1e). As expected, both Msd1-GFP and *γ*-Tubulin-GFP co-sedimented with MTs, demonstrating purified Augmin and *γ*-TuRC maintain at least some of their cellular properties.

To assess the effects of pure Augmin and *γ*-TuRC on MT nucleation and polymerisation, we used a highly-reproducible quantitative assay, where incorporation of a dye into MTs as they polymerise is measured as a change in fluorescence^26^ (Cytoskeleton Inc.). Incubation of Tubulin in the presence of GTP and glycerol at 37°C resulted in its polymerisation over ~1 hour, with sigmoidal dynamics corresponding to lag, nucleation, polymerisation and plateau phases (**Figure 2a**). The time at which 50% of polymerisation was achieved (x50) was 31.5mins (+/-0.5 mins) (**Figure 2b**). Addition of *γ*-TuRC-GFP stimulated MT nucleation, causing a shift in the polymerisation curve and a reduction in the x50 to 16.5 mins (+/-1.2 min) (**Figure 2a**,**b**), confirming its functionality. In contrast, addition of Augmin-GFP had no significant effect on the shape of the polymerisation curve or the x50 (32.5 mins (+/-1.5 min) (**Figure 2a**,**b**). Therefore, although Augmin-GFP binds MTs it does not, in isolation, change MT nucleation/polymerisation dynamics. However, addition of Augmin-GFP dramatically enhanced *γ*-TuRC-dependent nucleation of MTs, further reducing the x50 to 9.5 min (+/-0.45 min) (**Figure 2a**,**b**). This effect was specific for the physical interaction between Augmin and *γ*-TuRC, as addition of bacterially expressed and purified truncated Augmin subunits, Dgt3, Dgt5 and Dgt6, which we previously demonstrated interact directly with *γ*-TuRC^20^, resulted in nucleation/polymerisation curves indistinguishable to *γ*-TuRC alone (Supplementary Figure S2a-c). Thus, pure Augmin does, indeed, augment *γ*-TuRC-dependent MT nucleation *in vitro*.

**Figure 2.**
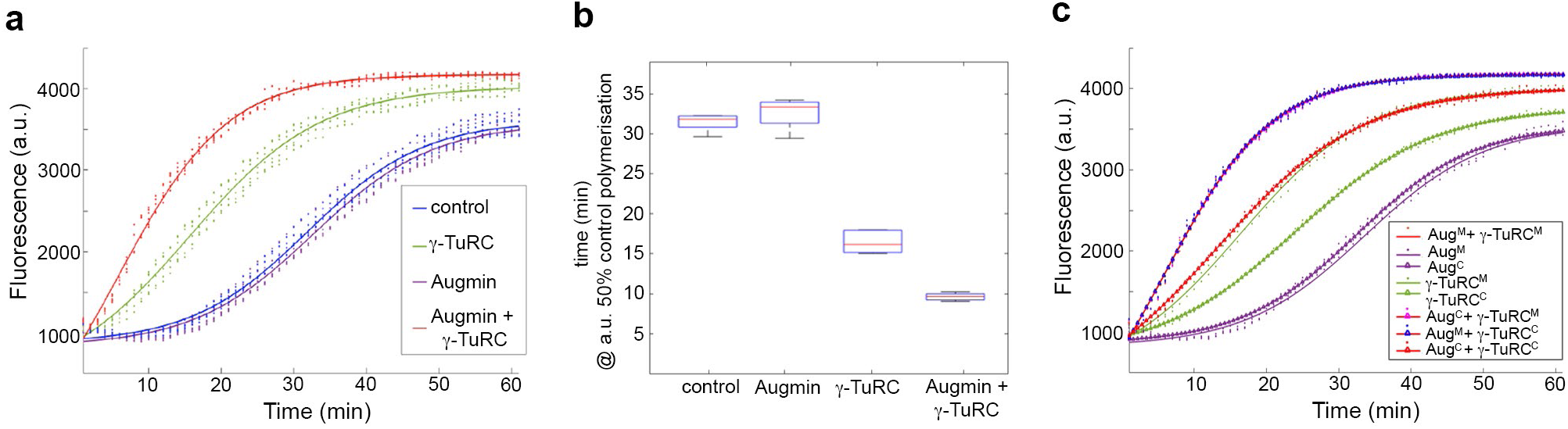
Pure Augmin enhances γ-TuRC-dependent MT nucleation in a cell cycle dependent manner. **a**. Tubulin polymerisation assays, where fluorescence is directly related to the amount of Tubulin polymer present. The curves are a sigmoidal fit to six data points (dots); three independent purification experiments, each undertaken in duplicate. **b**. plot of the x50 in relation to control polymerisation assay, showing the median (red line), interquartile ranges (blue box) and 95% confidence intervals of the median (notches) for each condition. The differences in the time taken for 50% polymerisation between all conditions differs significantly at p=0.001, except when comparing control to Augmin (ANOVA). **c**. fluorescent tubulin polymerisation assays undertaken with complexes isolated from MG132-treated (mitotic) or cycling embryos.

Importantly, this phenomenon was dependent on whether the complexes were isolated from mitotically arrested (MG132) or untreated, mainly interphase embryos (cycling). *γ*-TuRC purified from cycling embryos was a less efficient nucleator than its mitotic counterpart (**Figure 2c;** Supplementary Figure S3), while cycling Augmin, like mitotic Augmin, showed no independent activity. However, when incubated together, either mitotic *γ*-TuRC and cycling Augmin or cycling *γ*-TuRC and mitotic Augmin were able to enhance MT nucleation to the same extent as both mitotic complexes (**Figure 2c;** Supplementary Figure S3). As cell cycle dependent changes in protein function are determined mostly by post-translational modifications (PTMs), this observation suggests that PTMs of both Augmin and *γ*-Tubulin are crucial *in vivo*.

To characterise the morphology of MTs generated in the presence of pure Augmin and *γ*-TuRC, we took samples from the *in vitro* assays at t=15 minutes, fixing and imaging them via fluorescence microscopy. Control samples, or those containing pure Augmin, showed only very few, short MTs per field of view while, as expected, *γ*-TuRC-containing samples possessed many individual MTs (**Figure 3a**). In contrast, the MT population nucleated at t=15 simulataneously with *γ*-TuRC and Augmin showed a high density of MTs, with extensive MT bundling and nesting, rather than individual MTs (**Figure 3a**). Moreover, consistent with the observation that, *in vivo*, Augmin is required for *γ*-Tubulin localisation to the mitotic spindle, pure Augmin was able to recruit pure *γ*-TuRC to MTs *in vitro*, co-localising along the length of MTs in distinct punctae (Supplementary Figure S4).

**Figure 3.**
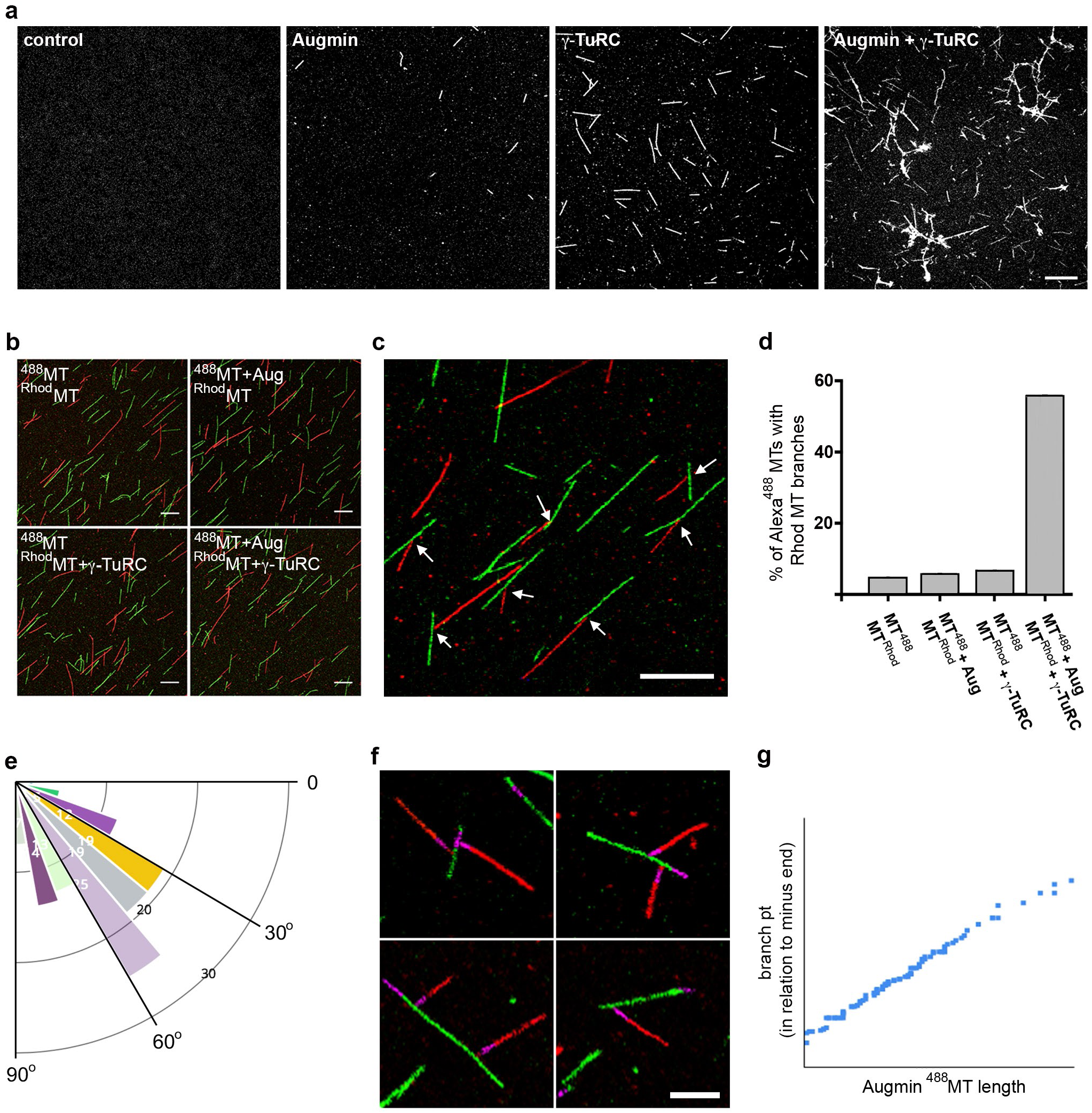
Reconstitution of the MT-Augmin-γ-TuRC-MT junction. **a**. confocal images of fixed, fluorescent polymerisation assays at t=15 mins for each condition. **b**. confocal images of taxol stabilised MTs, formed in the absence or presence of pure Augmin and γ-TuRC, and co-incubated. Incubation of Augmin-^488^MTs (green) with γ-TuRC-RhodMTs (red) leads to MT branches. c. higher magnification of the MT-Augmin-γ-TuRC-MT junctions (arrows). d. histogram of the percentage of ^488^MTs that possess a RhodMT branch. e. distribution of γ-TuRC-RhodMTs junction angles, in relation to Augmin-^488^MTs. **f**. confocal images of junctions, using GMPCPP seeds (purple) to distinguish the MT minus ends. Minus ends of γ-TuRC-RhodMTs (red) interact with Augmin-^488^MTs (green). **g**. Pearson correlation coefficient plot demonstrating a strong positive correlation between the length of Augmin-^488^MTs and the position of γ-TuRC-RhodMT branch points. Scale bars, c,e, 5µm; scale bar, f, 2µm.

Finally, we sought to conclusively reconstitute and visualise the branching MT-Augmin-*γ*-TuRC-MT junction. To overcome the complex morphology of MTs resulting from the nucleation assay, we generated populations of taxol-stabilised fluorescent MTs in the presence or absence of individual protein complexes, subsequently co-incubating them. Co-incubation of HiLyte Fluor 488-MTs (^488^MTs) with Rhodamine-MTs (^Rhod^MTs) resulted in independent populations of MTs. Occasional overlaps and apparent branching were observed, but at a frequency consistent with random placement of MTs on coverslips (**Figure 3b,d**). Similar results were obtained with fluorescent MTs incubated with either Augmin or *γ*-TuRC (**Figure 3b,d**). However, when ^488^MTs incubated with Augmin were added together with ^Rhod^MTs incubated with *γ*-TuRC, ~60% of *γ*-TuRC-containing ^Rhod^MTs terminated precisely at an Augmin-containing ^488^MT (**Figure 3b-d**). Although the polarity and angles of the ^Rhod^MTs branches in relation to the “mother” ^488^MTs varied far more widely than seen *in vivo*, presumably due to how the MTs settled on the coverslip during preparation, we found a bias in the angle, similar to that seen in living *Drosophila*^*16*^ (**Figure 3e**). To unequivocally demonstrate the polarity of the MTs, we formed these junctions in the presence of pre-stabilised HiLyte 647-labelled GMPCPP MT seeds, to additionally distinguish the slower-growing MT minus ends. As expected, the polarity of the interaction between *γ*-TuRC-^Rhod^MTs and Augmin-^488^MTs was specific; with only the minus ends of *γ*-TuRC-^Rhod^MTs precisely terminating at the Augmin-^488^MTs lateral surfaces (**Figure 3f**). Moreover, the formation of junctions on Augmin-containing ^488^MTs was length independent – branches were equally distributed at the very distal MT tips, throughout their length and on the GMPCPP seed itself (**Figure 3f,g**).

Our work confirms a long-standing hypothesis, first articulated to explain the loss of *γ*-Tubulin on the mitotic spindle when the expression of Augmin subunits is reduced^27^. It demonstrates conclusively that Augmin directly recruits *γ*-TuRC to MTs and that these two protein complexes are sufficient for the phenomenon of branched MT nucleation. It does not, however, exclude the possibility that *in vivo*, at least in some biological systems, other proteins tune branched MT nucleation to the individual needs of the cell. For example, a clear role has been reported for the MT associated protein, TPX2, in stimulating Augmin-dependent branched MT nucleation in *Xenopus* meiotic extracts^9,28^. However, in support of our conclusions, *Drosophila* TPX2 has recently been shown to be dispensible for the phenomenon *in vivo*^29^. Our experiments also highlight the intriguing possibility that, in some cells, Augmin might recruit pre-existing *γ*-TuRC-containing MTs, nucleated elsewhere in the cell, anchoring then to specific sites and increasing local MT density.

The generation of stable MT-Augmin-*γ*-TuRC-MT junctions using the methodologies pioneered here, also provide a route to finally defining the molecular detail of MT branching at the ultrastructural level. More broadly, cleavable affinity purification provides the basis to generate more complex, but molecularly defined, mixes of pure proteins, complete with *in vivo* PTMs, in order to reconstitute higher-order aspects of spindle formation. Indeed, by isolating and combining purified proteins and protein complexes from any biological system of interest, cl-AP overcomes the limitations of traditional “bottom-up” approaches, allowing exploration of the level of biological organisation between individual protein and biological process – the level at which emergence of cellular phenomena often occurs.

## Methods

### Drosophila husbandry and embryo collection

Flies were kept on standard medium and grown according to standard laboratory procedures. w1118 or Oregon R (OrR) strains (Bloomington Stock Center) were utilized as a wild-type controls. Transgenic flies used were: pUAS-Msd1-GFP; maternal-α-Tubulin-Gal4^26^ and pNcd(*γ*–Tub37C-GFP), a gift from Sharyn Endow^22^. 0-3-hour old embryos were collected from apple juice/Agar plates, dechorionated using bleach, washed, flash frozen in liquid nitrogen and stored at -80°C. For MG132 treatment, embryos were incubated in a solution containing 66.5% PBS (Melford), 33.2% heptane (Sigma) and 0.3% MG132 (Sigma) for 20 mins, prior to rinsing and flash freezing as above. Each embryo collection resulted in 0.05-0.1g of frozen embryos, different batches of which were combined for each biochemical assay/purification. Routinely, batches of MG132-treated embryos were fixed and stained to visualise chromosomes and MTs, to verify their mitotic arrest.

### Preparation of GFP-TRAP-PC and GFP-TRAP-Sulfo beads

To prepare GFP-TRAP-Sulfo beads, 200 μL of GFP-TRAP® (1 mg/mL) (Chromotek) in PBS was incubated with 86.1μl of Sulfo-NHS-SS-Biotin (sulfosuccinimidyl-20(biotinamido)ethyl-1,3-dithiopropionate) (8mM) (Thermo Scientific) for 2 hrs at 4°C. Unreacted linker was removed by desalting. 40 μL of High Capacity Streptavidin Agarose Resin (Pierce) was washed 3 times for 5 mins in PBS and incubated with the GFP-TRAP-SS-Biotin product for 1 hr at 4°C with gentle rotation. The beads were washed 3 times for 5 mins in BRB80 + 0.1% IGEPAL®, the volume of BRB80 was reduced to 40 μL (to maintain a 50% bead slurry) and beads stored at 4°C for use. A similar protocol was used to generate GFP-TRAP-PC-Biotin-NHS beads, but using 50 μL of GFP-TRAP®, incubated with 1.45 µL of PC-Biotin-NHS (50mM) (Ambergen) for 40 minutes at RT with gentle rotation and 400 µL of standard Streptavidin-Agarose bead slurry (Sigma Aldrich). Over the course of the project, different conditions were tried in an attempt to increase the photo-cleaving of GFP, GFP-Augmin and GFP-*γ*-TuRC from the GFP-TRAP-PC beads. We were unable to consistently reach cleaving efficiencies of greater than 60%. In contrast, cleavage of proteins and complexes from GFP-TRAP-Sulfo-beads reproducibly gave efficiencies of >90% (n=>10).

### Cleavable Affinity Purification

*Drosophila* embryo High Speed Supernatant (HSS) was prepared using batches of frozen 0-3 hr embryos. Embryos were dounce homogenised in BRB80 + 0.1% IGEPAL + 1mM PMSF (Sigma), PhosSTOP phosphotase inhibitors and cOmplete, Mini, EDTA free protease inhibitors (Roche) at a ratio of 100mg embryos to 200 μL buffer. Extracts were clarified through centrifugation at: 17,000 g for 10 min, 100,000 g for 10 min, and 100,000 g for a further 30 min. 6xHis-GFP was purified from bacteria expressing pQE80-His-GFP, using standard protocols and HisPur™ Cobalt Resin (Thermo Scientific), and dialysed into BRB80 + 0.1% IGEPAL + 1mM PMSF.

Typically, HSS from 200 mg embryos or 6xHis-GFP (100ng) were incubated overnight at 4°C with 20 μL of either GFP-TRAP-PC or GFP-TRAP-Sulfo beads. For SYPRO-Ruby staining (see below) HSS from 1 g of embryos expressing either Msd1-GFP or *γ*-Tubulin-GFP were used, incubated with 50 μL of GFP-TRAP-Sulfo beads. Beads were centrifuged at 2500 g for 2 mins and the depleted supernatant removed and discarded, or used for Western blotting. For cleaving of Sulfo-beads, beads were washed 4 times for 5 mins in BRB80 + 0.1% IGEPAL®, resuspended in 25 μL of BRB80 + 0.1% IGEPAL®, prior to addition of 25 μL of 100 mM DTT in BRB80 0.1% IGEPAL® for 5 mins (50mM final concentration). For PC-beads, beads were washed as above, then transferred to a microscope slide with a concave cavity with 25 μL of BRB80 0.1% IGEPAL®. The slide was placed 10cm below a UV lamp (UVP XX-15L (Analytik Jena US)) and exposed for 30 second intervals with gentle mixing between. Eluates were removed using a Gel-Saver II pipette tip (STARLAB). Cleaved beads were washed 4 times for 5 minutes in BRB80 0.1% IGEPAL®. After each purification, samples were taken for SDS-PAGE and Western blotting alongside known amounts of 6xHis-GFP, in order to quantify yield. cl-AP of 200 μg of embryos using GFP-TRAP-Sulfo beads consistently yielded ~400 ng of Msd1-GFP and *γ*-Tubulin-GFP (50 μl at 8-9 ng/uL) (n=6).

### SDS-PAGE and Western blot analysis

Protein samples were fractionated by sodium dodecyl sulfate (SDS)-polyacrylamide gel electrophoresis (PAGE). SYPRO Ruby Staining (Invitrogen) of Augmin and *γ*-TuRC was undertaken according to manufacturer’s instructions. For Western analysis, the proteins were blotted onto a nitrocellulose membrane and probed with anti-GFP (1:1000; Sigma-Aldrich and Roche), anti-dgt6 (1:500) (a gift from M. Gatti), or anti-Dgrip91 antibodies (1:1000) (a gift from X. Zheng). IRDye 800CW goat anti-rabbit (LI-COR) and IRDye 680RD goat anti-mouse (LI-COR) IgG polyclonal Abs were used as secondary detection antibodies. Fluorescence from blots was developed with the Odyssey CLx Imaging System (LI-COR) according to the manufacturer’s instructions.

### *In Vitro* Microtubule Co-sedimentation Assay

Purified GFP, GFP-Augmin or GFP-*γ*-TuRC were pre-spun at 100,000 g for 15 min at 4°C. 10 μL (80 ng) of each was incubated for 15 min at 37°C in General Tubulin Buffer with 2.25 mg/mL of 99% pure porcine tubulin and 1 mM GTP (all from Cytoskeleton Inc.), in a final volume of 50 μL. Taxol (Sigma) was added to 100 µM and samples incubated for a further 10 min at 37°C. A negative control for each sample was run in parallel, where samples were incubated at 4°C and where GTB replaced taxol. Samples were immediately spun through a 150 µL cushion of BRB80 40% glycerol at 100,000 g for 45 min at 4°C in a TLA120.1 rotor (Beckmann Coulter). The supernatant and pellet fractions were analysed by Western blotting, probing with mouse anti-GFP (Roche) and anti-*α*-Tubulin antibodies [DM1A] (Sigma). Co-sedimentation assays were undertaken in triplicate, each with independently purified Augmin or *γ*-TuRC. A representative western is shown.

### MT polymerization assays

MT polymerization assays were performed using a fluorescence-based Tubulin Polymerization kit (Cytoskeleton Inc, Denver CO, Cat. # BK011P) following the manufacturer’s instructions. Briefly, 5 μL (40 ng) of GFP, Augmin or *γ*-TuRC was pipetted into wells within a 96-well microtiter plate, followed by 45 μL of Tubulin Reaction Mix. Tubulin polymerization was initiated by transferring the plate to a 37°C chamber of a plate reader. The polymerization dynamics of Tubulin were monitored for 60 min at 37°C by measuring the change in fluorescence every 1 min using a TECAN infinite 200pro fluorimeter, at excitation of 350 nm and emission of 440 nm. Assays to assess the effect of pure Augmin and *γ*-TuRC presented in Figure 2A are the summation of 3 individual purifications of each protein complex, undertaken in duplicate wells (6 data points). Assays to assess the difference in polymerization between cycling and MG132 purified Augmin and *γ*-TuRC, presented in Figure 2C are the summation of triplicate experiments. *In vitro* polymerization assays to assess the difference in Augmin-dependent polymerization upon addition of competing truncated Augmin subunits (Supplementary Figure S2) are the summation of triplicate experiments.

### Generation of fluorescent MTs

Tubulin Polymerization assays were performed as described above, but with the following modifications: Rhodamine- or HiLyte Fluor 488-tubulin was used (Cytoskeleton, Inc., Denver, CO) at a 1:10 ratio with unlabelled porcine tubulin (final tubulin concentration, 2mg/ml). At t=15 min, samples of MTs were incubated with glutaraldehyde (0.5 % final concentration) for 15 mins. Fixed MTs were spotted onto glass coverslips precoated with mounting media (85% glycerol; 2.5% N,N propyl-gallate) and imaged.

To generate fluorescent, taxol stabilised MTs, Rhodamine- or HiLyte Fluor 488-tubulin was polymerized as above, but in the absence of pure complexes, but in the presence of taxol at a final concentration of 20 μM. After 7 mins, the taxol-stabilized MTs were removed from the well and incubated with 5 μL (40 ng) of Augmin, *γ*-TuRC or buffer for 10 min at room temperature. Samples were then layered over a 150 μL cushion of 15% glycerol in BRB80 and centrifuged at 135,000 ×g for 10 mins at 25° C in a TLA120.1 rotor (Beckmann Coulter). The MT pellets were gently resuspended in 50 μL of General Tubulin Buffer containing 1mM GTP and 30% glycerol, combined as necessary as 1:1 mixtures and incubated for 10 min at room temperature. The samples were fixed with glutaraldehyde (0.5-2 % final concentration) and spotted onto glass coverslips as above. Images were taken from randomly distributed fields of the coverslips using a Leica TCS SP8 confocal laser scanning microscope. Representative images from one of three independent experiments are shown.

For experiments to visualise MT polarity, HiLyte 647-labelled GMPCPP MTs were generated according to the manufacturer’s instructions (Jena Biosciences). Seeds were added to the polymerisation reactions at a 1:10 ratio and processed as described above. Images were taken from randomly distributed fields of the coverslips using a Leica TCS SP8 confocal laser scanning microscope. Representative images from one of two independent experiments are shown.

### Imaging and Image analysis

Samples were imaged using an inverted Leica TCS SP8 confocal laser scanning microscope using a HCOL APO CS2 63x, NA 1.4 oil immersion lens (Leica, Wetzlar, Germany). Standard filter sets were used to visualize Rhodamine, HiLyte Fluor 488, and HiLyte Fluor 647 fluorescence. Images were captured as TIFFs using Leica Application Suite X (LAS X), opened in FIJI and levels adjusted to maximise the full range of pixel intensities. To quantify the % of ^488^MTs with ^Rhod^MT branches, 8 fields of view of each sample were randomly captured. ^488^MTs over ~1μM length in each field of view were totaled, alongside the number of ^Rhod^MTs whose fluorescence terminated precisely at a HiLyte^488^MT lateral surface (between 243 – 339 ^488^MTs per sample condition). To quantify the position of the branchpoints relative to the minus ends of MTs, the length of HiLyte^647^ GMPCPP-containing ^488^MTs were measured using the line tool in ImageJ. The position of the ^Rhod^MT branch from the minus end of these MTs, was also measured. The two sets of variables were subjected to Pearson correlation coefficent analysis using an on-line tool (https://www.socscistatistics.com/tests/pearson/default2.aspx), <u>providing the graph in</u> Supplementary Figure 3. The Pearson correlation coefficient was r=0.9962 at a p value of <0.00001.

### Statistical analysis of polymerisation curves

Polymerisation data sets were fitted using a sigmoidal function. This characterised each data set in terms of the maximum & minimum fluorescence, the x value (min) at 50 % distance between maximum and minimum y fluorescence value (termed ‘x50’) and the slope. The mean x50 value for the control samples (n=6) was x=31.99 (minutes) and y=2411 (fluorescence a.u.). This value was used as a fix point to compare all the data sets. A one-way ANOVA was performed to compare the four variables (control, Augmin, *γ*-TuRC, and Augmin+*γ*-TuRC), with each variable having six repeats. A *post-hoc* Tukey test showed that all variables were significantly different from each other at *p*=0.001 apart from the control and Augmin, which were not significantly different from each other. Similar analyses were undertaken for the comparison of polymerisation datasets between complexes isolated from MG132 and cycling embryos, and the augmenting activity of Augmin in the presence of truncated Dgt3, Dgt5 and Dgt6 proteins. All fitting and analyses were performed in MATLAB (the code and data can be found at www.github.com/charliejeynes/microtubles).

## Acknowledgements

We thank Stephen Green and Mark Wood (Exeter, UK) for initial discussions about photocleavable and other tags, and Mary Munson (Massachusetts, US) for highlighting the potential of thiol-cleavable tags for native complex purification, as published by the Rout lab (Rockefeller, US)^30^. We thank Chromotek for the gift of GFP-TRAP; Jack Chen, Dan Hayward and Chris Sullivan for initial attempts to develop the cl-AP technique; Marwan Al-Maqtoofi who supervised a set of University of Exeter Natural Sciences undergraduate students and the Carlota Palmer PhD students in setting up the plate reader-based MT nucleation/polymerisation assays. Finally, we thank Carolyn Moores (Birkbeck, UK), Iris Leuke and Thomas Surrey (Crick Institute, UK) for discussions and advice, and Sabine Petry for sharing unpublished data and critical reading of the manuscript. L.G. was supported by a University of Exeter Proof of Concept Award, A.T. by a Carlota Palmer University of Exeter PhD Studentship and J.C.G.J. by an Innovation Fellowship supported by the Science and Technology Facilities Council (STFC), U.K.

## Author contributions

J.G.W. designed the study. L.G. developed the GFP-TRAP-PC approach and undertook associated experiments. A.T. conducted all other experiments. J.C.G.J. analysed the polymerisation assay data. C.S. co-supervised A.T. co-designed and analysed the imaging experiments. J.G.W. and A.T. analysed all other data and wrote the manuscript.

