## Supplemental Figure S1 for "*In vitro* reconstitution of branched microtubule nucleation"

### Supplementary Figure S1

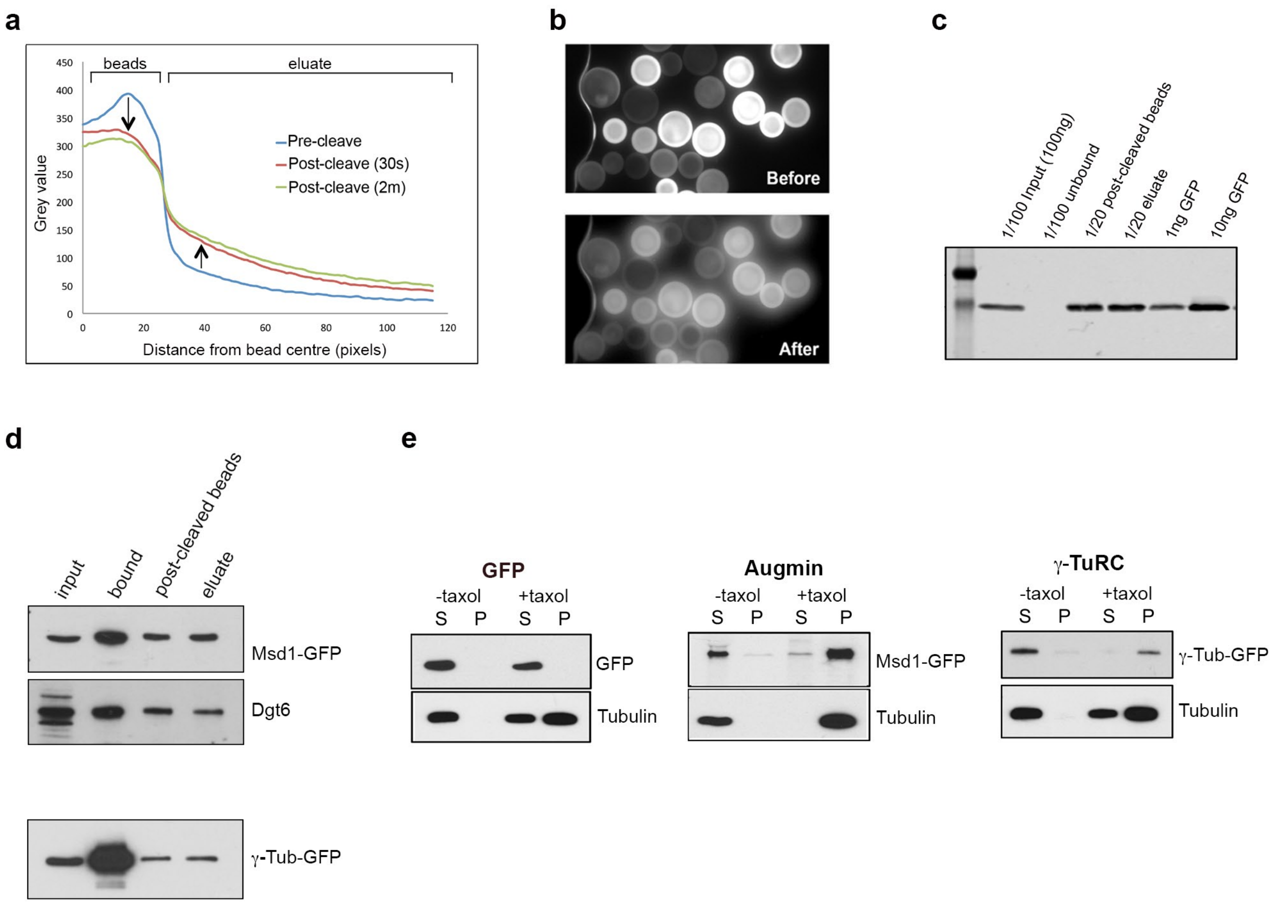


**Supplementary Figure S1** Isolation of functional γ-TuRC and Augmin using GFP-TRAP-PC beads. **a.** Graph showing the comparative decrease in GFP fluorescence on GFP-TRAP-PC beads, and increase in the surrounding eluate, following exposure to UV light (UV filter block on TE2000U fluorescence microscope). **b.** Images of GFP-TRAP-PC beads after incubation with GFP, pre- and post-cleavage by 30s exposure to UV light, used to generate data for (a). **c.** Western blot to quantify the binding and release capacity of GFP-TRAP-PC beads. 100ng of GFP is completely immobilised on the beads. In this case, ~60% of GFP is cleaved via UV exposure, corresponding to 3ng x 20 = 60ng GFP. d. Western blots demonstrating the isolation and cleavage of Msd1-GFP and γ-Tubulin-GFP using GFP-TRAP-PC beads. The proteins, present in embryo extracts (input), are efficiently captured onto beads (bound), with ~50% released following UV exposure (post-cleaved beads and eluate). **d.** Western blots of *in vitro* MT co-sedimentation assays using GFP-TRAP-PC beads. In the absence of taxol (-), Tubulin, pure Augmin (Msd1-GFP) and pure γ-TuRC (γ-Tubulin-GFP) remain in the supernatant (S). In the presence of taxol (+), Tubulin polymerises and is present in the pellet (P). Augmin and γ-TuRC co-sediment. GFP alone does not co-sediment with MTs in this assay.
