## Supplemental Figure S2 for "*In vitro* reconstitution of branched microtubule nucleation"

### Supplementary Figure S2

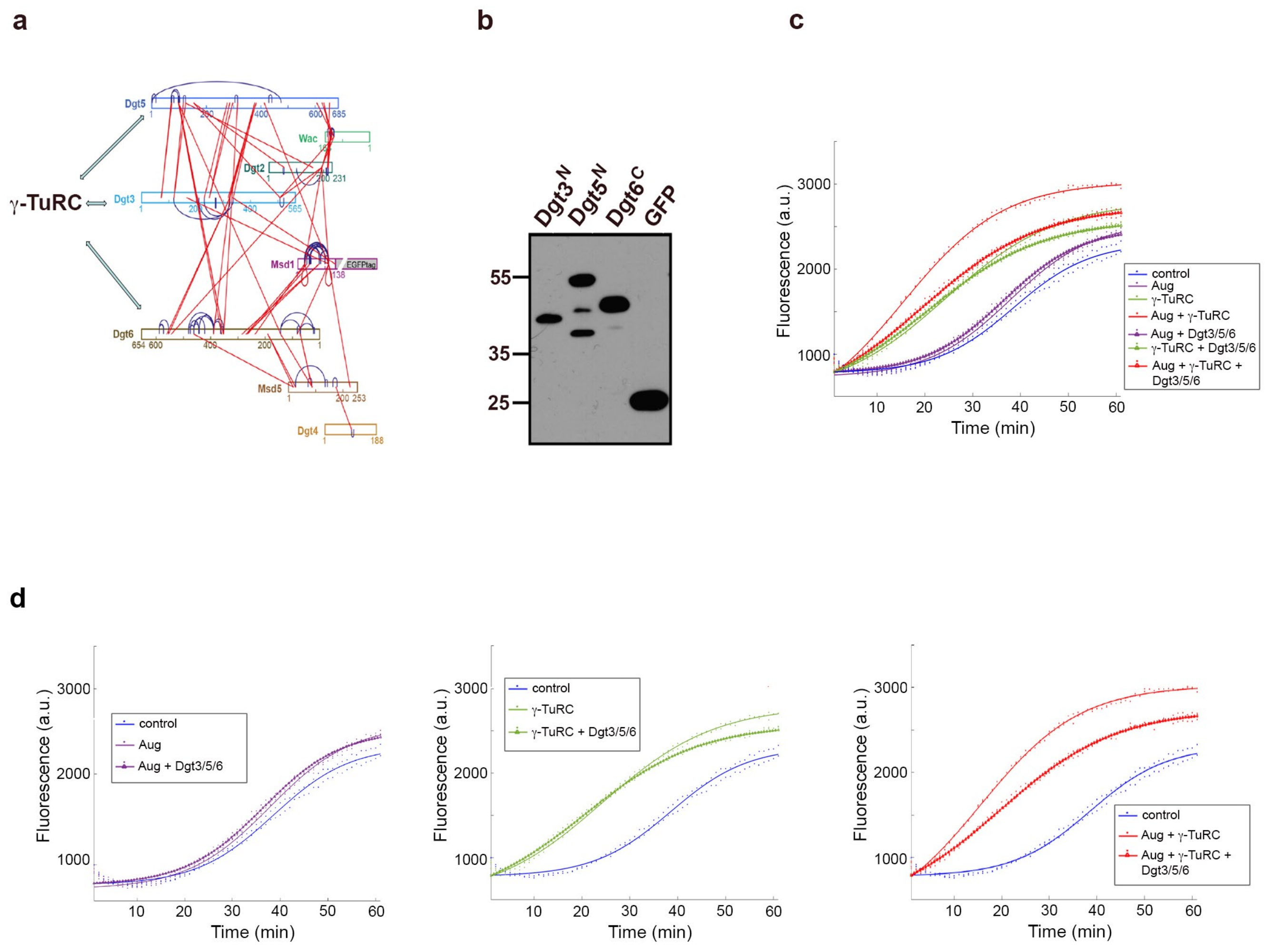


**Supplementary Figure S2** The synergistic effect of pure Augmin on γ-TuRC-dependent MT nucleation *in vitro* is dependent on the Augmin-γ-TuRC interface. **a.** representation of the interactions between Augmin subunits, as determined by cross-linking mass spectrometry. The N-terminus of Dgt3 and Dgt5, and the C terminus of Dgt6 have all been previously shown to be responsible for localising the γ-TuRC to the mitotic spindle in *Drosophila* embryos (from Chen et al., 2017). **b.** western blot showing the purification of MBP-Dgt3$^{N}$, MBP-Dgt5$^{N}$ and MBP-Dgt6$^{C}$, expressed in bacteria (adapted from Chen et al., 2017). **c.** fluorescent tubulin polymerisation assays undertaken with GFP-Augmin and GFP-γ-TuRC isolated from MG132-treated embryos, in the presence and absence of Dgt3$^{N}$, Dgt5$^{N}$ and Dgt6$^{C}$. Inclusion of the Dgt regions completely abrogates the ability of Augmin to affect γ-TuRC-dependent MT nucleation, without affecting γ-TuRC-dependent MT nucleation itself. d. Individual datasets from (c). Note that inclusion of the Dgt regions with γ-TuRC alone does not affect γ-TuRC-dependent MT nucleation but does slightly reduce the total amount of MT polymer formed (middle graph).
