## Supplemental Figure S3 for "*In vitro* reconstitution of branched microtubule nucleation"

### Supplementary Figure S3

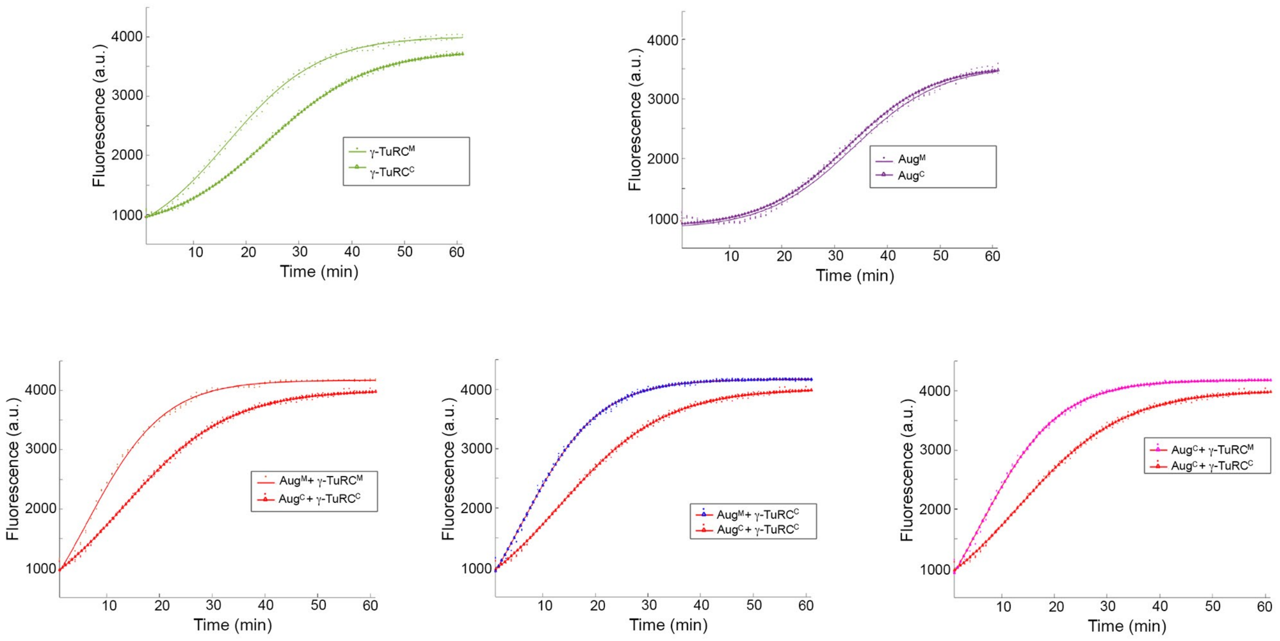


**Supplementary Figure S3** Individual sets of polymerisation curves using Augmin and γ-TuRC purified from either mitotic (M) or cycling (C) embryo extracts, taken from Figure 2c. Mitotic γ-TuRC stimulates MT nucleation/polymerisation to a greater extent than γ-TuRC isolated from cycling embryo extracts. Neither cycling nor mitotic Augmin affects MT nucleation/polymerisation. Addition of mitotic Augmin to mitotic γ-TuRC stimulates γ-TuRC-dependent MT nucleation. This effect is reduced when cycling Augmin and cycling γ-TuRC are used. However, if either Augmin or γ-TuRC is mitotic, Augmin-dependent MT nucleation is active, even if the other complex is isolated from cycling embryos.
