## Supplemental Data 1 for "*In vitro* reconstitution of branched microtubule nucleation"

### Supplementary Figure S4

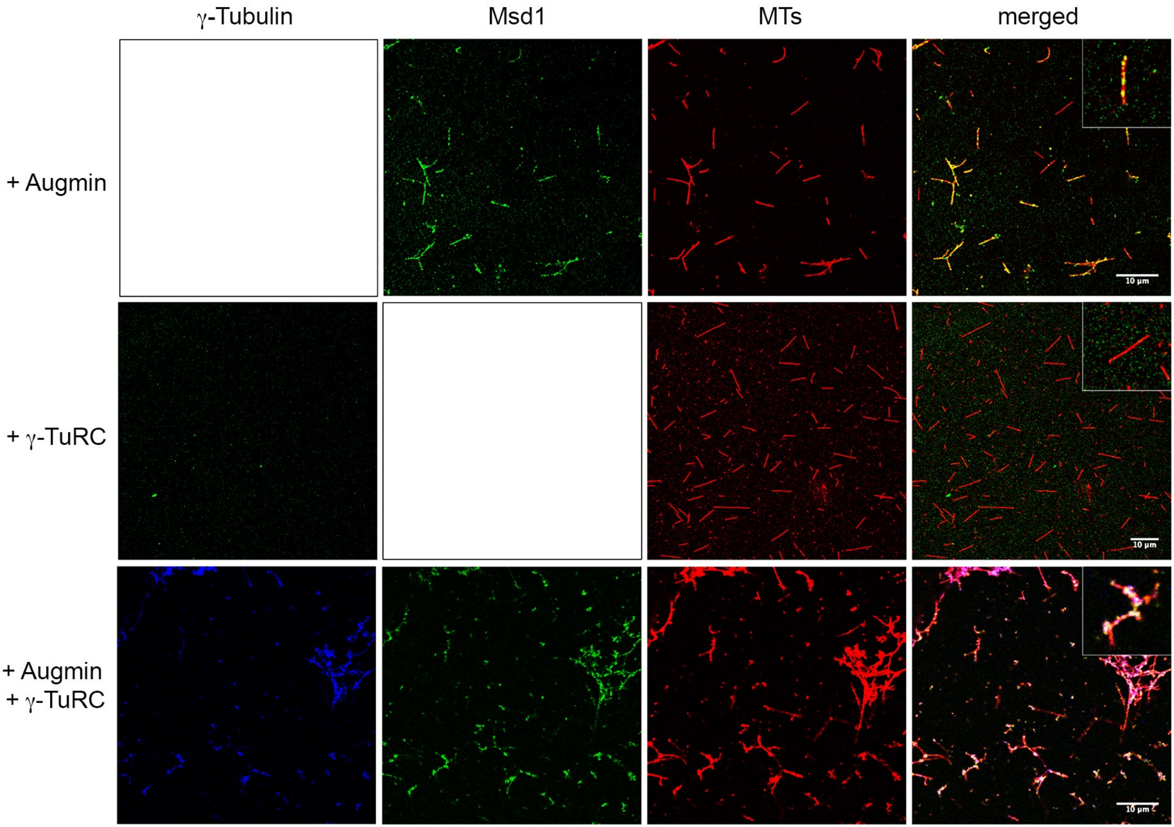


**Supplementary Figure S4** Augmin recruits γ-TuRC to MTs. Samples were taken mid-way through in vitro polymerisation reactions, fixed and stained for γ-Tubulin, the Augmin Msd1 subunit or MTs. When present on its own, Augmin localises along the length of MTs. Note this localisation is not uniform; rather Augmin is present as punctae. When γ-TuRC is present on its own, γ-Tubulin is not observed on MTs, even though it stimulates MT nucleation (presumably the levels on the minus ends of MTs are too low to be resolved). However, when γ-TuRC is present with Augmin, γ-Tubulin co-locaises with Augmin on MTs. Scale bar, 10 μm. See insets for higher magnified images.
